# Deep self-supervised learning for biosynthetic gene cluster detection and product classification

**DOI:** 10.1101/2022.07.22.500861

**Authors:** Carolina Rios-Martinez, Nicholas Bhattacharya, Ava P. Amini, Lorin Crawford, Kevin K. Yang

## Abstract

Natural products are chemical compounds that form the basis of many therapeutics used in the pharmaceutical industry. In microbes, natural products are synthesized by groups of colocalized genes called biosynthetic gene clusters (BGCs). With advances in high-throughput sequencing, there has been an increase of complete microbial isolate genomes and metagenomes, from which a vast number of BGCs are undiscovered. Here, we introduce a self-supervised learning approach designed to identify and characterize BGCs from such data. To do this, we represent BGCs as chains of functional protein domains and train a masked language model on these domains. We assess the ability of our approach to detect BGCs and characterize BGC properties in bacterial genomes. We also demonstrate that our model can learn meaningful representations of BGCs and their constituent domains, detect BGCs in microbial genomes, and predict BGC product classes. These results highlight self-supervised neural networks as a promising framework for improving BGC prediction and classification.

**Author summary:** Biosynthetic gene clusters (BGCs) encode for natural products of diverse chemical structures and function, but they are often difficult to discover and characterize. Many bioinformatic and deep learning approaches have leveraged the abundance of genomic data to recognize BGCs in bacterial genomes. However, the characterization of BGC properties remains the main bottleneck in identifying novel BGCs and their natural products. In this paper, we present a self-supervised masked language model that learns meaningful representations of BGCs with improved downstream detection and classification.

## Introduction

Natural products are chemical compounds that form the basis of many pharmaceuticals and clinical therapeutics [1]. Their chemical structures are used in the development of antimicrobial drugs, anticancer therapies, and other therapeutic areas [2]. To initiate the discovery of natural products, the pharmaceutical industry has traditionally relied on laboratory research, yet this approach cannot feasibly capture the entire chemical diversity of natural products. Thus, new methods are needed to advance natural product discovery [3].

Diverse natural products can be produced in living organisms via groups of genes called biosynthetic gene clusters (BGCs). Genome mining has become a powerful tool for exploring the complex and diverse chemical space of natural products [3]. Fast, inexpensive genome sequencing technology has contributed to the advancement of BGC identification and, by extension, natural product discovery. This approach has been particularly successful in microbes, where BGCs are often a group of physically colocalized genes whose sequence and function dictates the synthesis of natural products. However, evidence suggests that much of the biosynthetic capacity of the microbial world remains unexplored [4]. Improved identification and characterization of BGCs directly from genomic data could accelerate the discovery of novel natural products with therapeutic relevance.

Identification of BGCs directly from genomic sequences is critical to navigating natural product space and nominating novel natural products. While complementary data modalities involving joint genome sequencing and mass-spectrometry data can be used to link products with gene clusters [5], the majority of known BGCs were characterized directly from DNA sequencing performed without any associated analysis of chemical structures in the sample. As such, computational methods which focus exclusively on identifying BGCs from genomes are essential components of BGC discovery pipelines.

antiSMASH (ANTIbiotics & Secondary Metabolite Analysis SHell) is an early tool for BGC discovery that uses a set of curated profile-Hidden Markov Models (pHMMs) to call biosynthetic gene families and a set of heuristics to tag a portion of a genome as a BGC [6, 7]. antiSMASH then annotates these called BGCs by using carefully curated rules based on expert knowledge. Similarly, ClusterFinder uses a Hidden Markov Model (HMM) to identify gene clusters of known and unknown classes [8]. Despite their effectiveness, HMM-based algorithms do not capture higher-order dependencies between genes, limiting their accuracy and generalizability [9]. Likewise, rule-based methods are limited by the need for human expertise and do not generalize well to new BGC classes.

A recent approach, DeepBGC, introduced a deep learning genome-mining strategy for biosynthetic gene cluster annotation that addresses these limitations [10]. Similar to antiSMASH, DeepBGC uses sets of curated pHMMs to call biosynthetic gene families; however, it uses a supervised neural network to predict BGC boundaries and annotate BGC function. Specifically, they employ a bidirectional long short-term memory (Bi-LSTM) recurrent neural network (RNN), which offers the advantage of capturing short- and long-term dependencies between adjacent and distant genes [11]. DeepBGC reported promising improvements in the identification of BGCs in microbial genomes. However, DeepBGC is trained on a small number of high-quality annotations, and the supervised approach requires mining examples of genes that are not part of BGCs. The quality of the predictions is highly dependent on the quality of the negative examples, which must be similar to BGC sequences while ideally containing no false negatives.

Rather than relying on expert-curated annotations and negative examples, self-supervised masked language models promise the ability to learn biologically-relevant patterns directly from a large set of BGC examples. Recently, self-supervised masked language models of biological sequences have been used to study proteins [12–19], DNA [20], RNA [21, 22], and glycans [23, 24]. In these models, a neural network is trained either to reconstruct the original sequence from a corrupted version of the sequence, or to predict the next element in the sequence given the preceding elements. After training on a large dataset, such as all protein sequences in UniProt [25], the model can be used for zero-shot predictions of fitness [26] or structure [27], and can additionally be fine-tuned on downstream supervised tasks [28, 29].

To accelerate identification and classification of BGCs, we developed a self-supervised neural network masked language model of BGCs from bacterial genomes (Fig. 1). Our model represents BGCs as chains of functional protein domains, and uses ESM-1b [12], a protein masked language model, to obtain pretrained embeddings of functional protein domains with amino acid-level context. We then train a convolutional masked language model on these domains to develop meaningful learned representations of BGCs and their constituent domains. The architecture for our model is based off of convolutional autoencoding representations of proteins (CARP) [30], a masked language model of proteins, and we will therefore refer to it as **Bi**osynthetic **G**ene **CARP** (BiGCARP). We leverage these representations to detect BGCs from microbial genomes and then classify them based on their natural product class. We further investigate the potential advantages of our model by comparing our approach with DeepBGC, and demonstrate that BiGCARP achieves improvements in BGC prediction and natural product classification. BiGCARP highlights self-supervised neural networks as a promising framework for improving BGC characterization.

**Fig 1.**
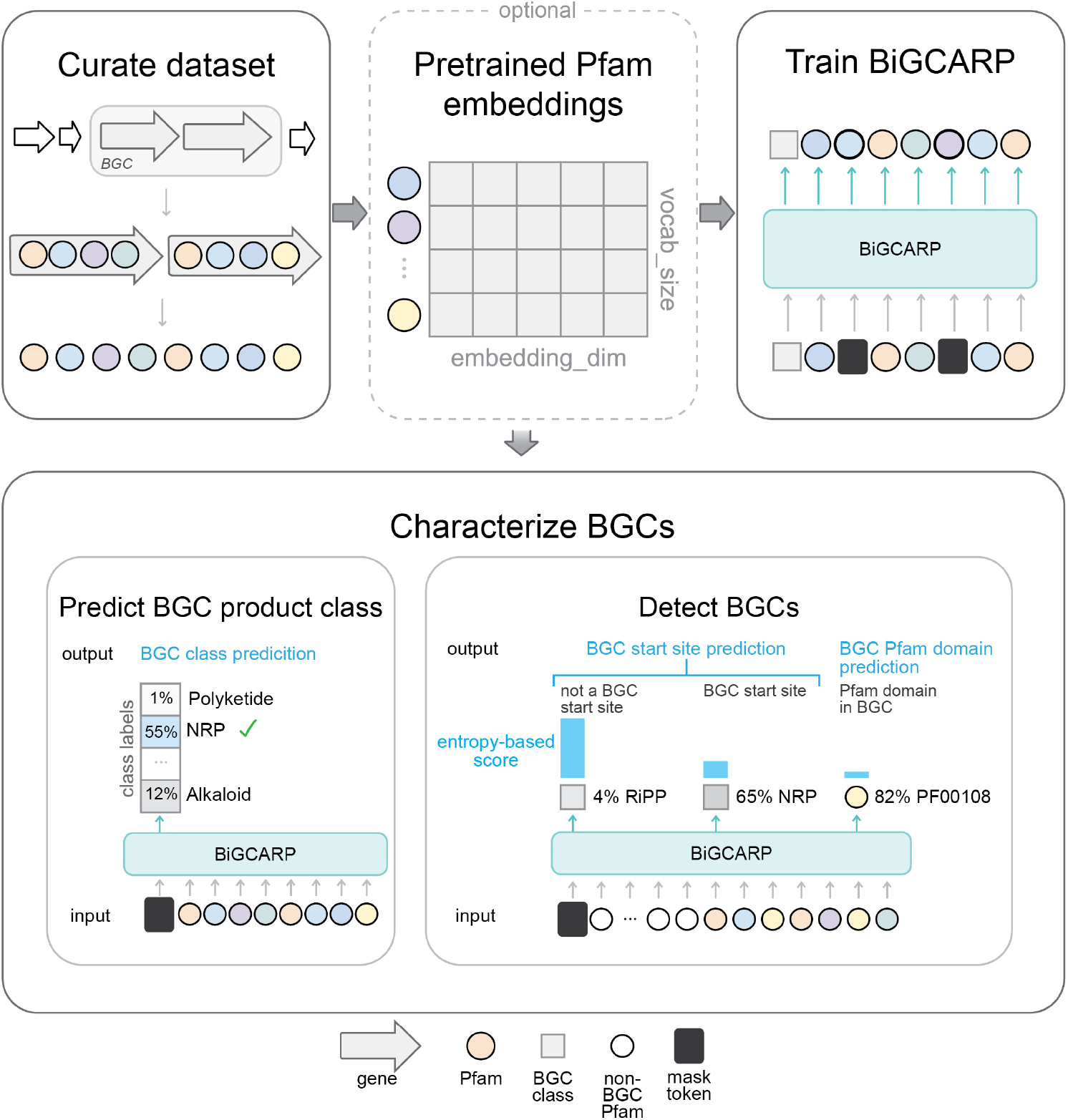
Self-supervised deep learning workflow for characterizing biosynthetic gene cluster (BGC) properties. Schematic of the workflow for characterizing BGCs with BiGCARP, a self-supervised deep neural network. We curate a dataset of annotated BGCs from antiSMASH for training BiGCARP. We then use ESM-1b [12], a protein masked language model, to obtain pretrained embeddings of protein family (Pfam) domains in our dataset and to explore whether pretrained Pfam domain embeddings show improvement on the quality of their representations. By representing BGCs as chains of Pfams, we train a self-supervised masked language model on these domains to characterize BGC properties in microbial genomes. We leverage these learned representations to detect BGCs from microbial genomes and to predict their natural product class.

## Results

### Self-supervised training

We first developed a self-supervised training scheme to train BiGCARP to learn representations of BGCs. As BGCs have a hierarchical structure, they can be represented at four main levels. From least-to-most granular, these are: genes, Pfam domains (families of evolutionary-related proteins), amino acids, and nucleotides. We note that more granular units of representation lead to longer sequences. BGCs typically contain several dozen genes, each of which contains one or more Pfam domains. Each Pfam domain contains tens to hundreds of amino acids, and each amino acid is encoded by three nucleotides. This introduces a trade-off between modeling short sequences where each unit is complex or modeling long sequences where each unit is simple. In order to balance input sequence length and information content of individual units, we chose to represent BGCs as sequences of Pfams. This is the same level chosen by DeepBGC [10]. As shown in Fig. 1, during training, we append a BGC product class token to the start of each BGC Pfam sequence in order to learn BGC product classes from their Pfam domain sequences. We then corrupt the sequence according to the BERT [31] corruption scheme and train **Bi**osynthetic **G**ene **C**onvolutional Autoencoding **R**epresentations of **P**roteins (BiGCARP) to reconstruct the original class token and Pfam sequence. BiGCARP combines the ByteNet encoder dilated CNN architecture from [32] with linear input embedding and output decoding layers, as shown in Figure 2a.

**Fig 2.**
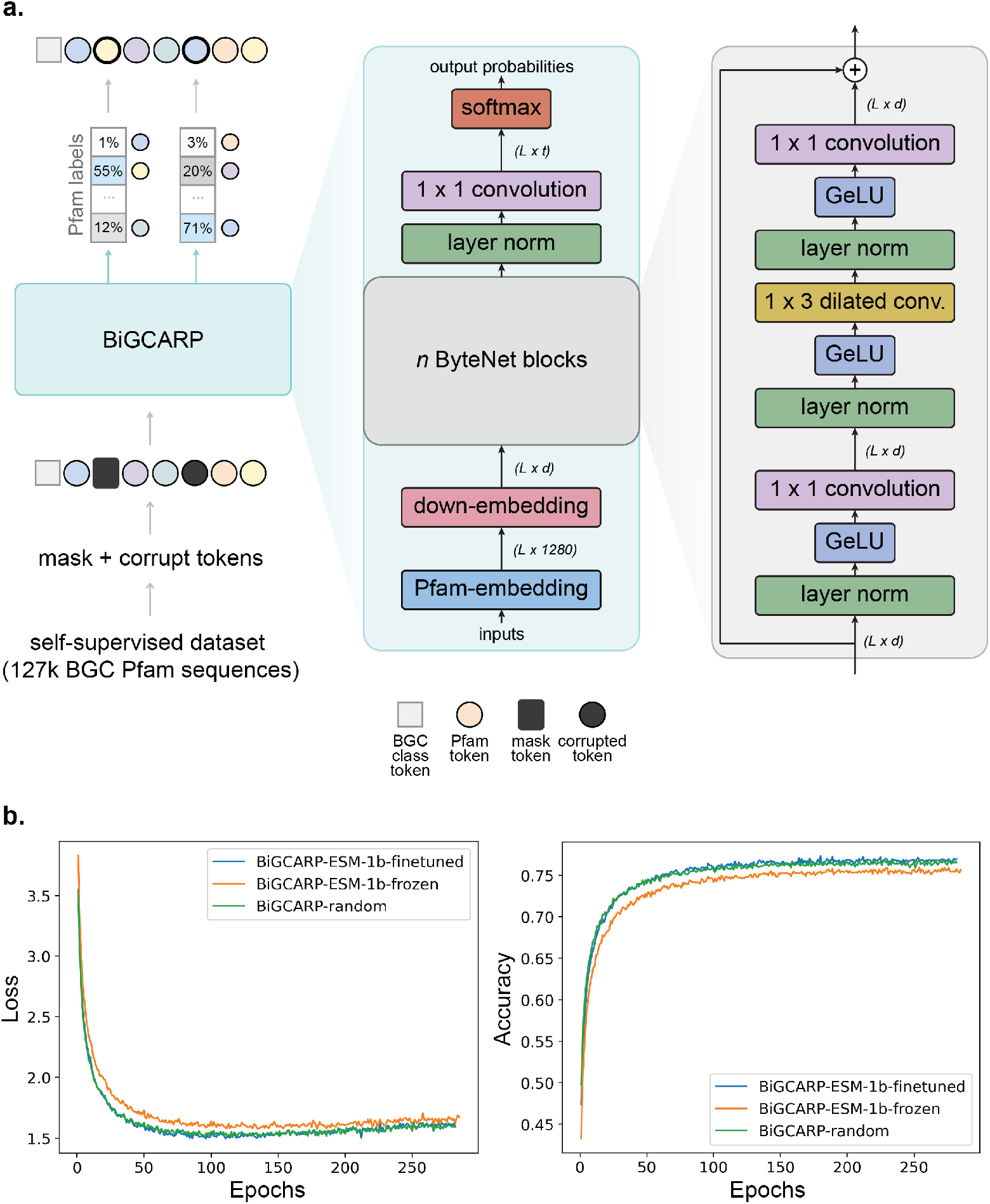
BiGCARP architecture with validation performance curves on the self-supervised dataset. (a) We use the masked language model objective described in [31] to train BiGCARP to reconstruct the BGC product class and Pfam sequence on our self-supervised dataset, which contains around 127,000 BGC Pfam sequences. BiGCARP is a dilated 1D-convolutional neural network masked language model based on CARP [30] and ByteNet [32]. (b) Validation loss (cross-entropy) and accuracy for BiGCARP with different initial Pfam embeddings.

Pfam embeddings map protein families from our vocabulary to vectors in a high-dimensional space, and thus serve as the inputs to BiGCARP. We train three versions of BiGCARP with different initial Pfam embeddings. The BiGCARP-ESM-1b-finetuned and BiGCARP-ESM-1b-frozen models are both initialized with Pfam embeddings obtained by averaging the per-residue output from ESM-1b for each domain. BiGCARP-ESM-1b-finetuned has its embeddings finetuned during self-supervised BGC training, while BiGCARP-ESM-1b-frozen has the initial embeddings frozen at the onset of self-supervised BGC training. Finally, BiGCARP-random is initialized with a random Pfam embedding, which is finetuned during self-supervised BGC training. All three versions of BiGCARP are trained on BGC sequences extracted from the antiSMASH dataset [6, 7]. We used approximately 127,000 BGC sequences and split the dataset 80/10/10 between training, validation, and testing, respectively. The training set is deduplicated against all datasets used in downstream evaluation. We refer the reader to Materials and Methods for details about the model training and architecture and the self-supervised training dataset.

Figure 2b plots the learning curves of the validation performance on the self-supervised dataset for all three versions of BiGCARP. We discover BiGCARP-ESM-1b-frozen is outperformed by BiGCARP-ESM-1b-finetuned and BiGCARP-random, which both show similar performance and attain an accuracy of around 75%.

### Learned embeddings encode relevant representations of Pfam domains

We used uniform manifold approximation and projection (UMAP) to visualize the input Pfam embeddings after self-supervised training on the antiSMASH training set (Fig. 3). Each protein family is represented as a single point, and protein families of similar sequence and function should have similar representations and thus be mapped to nearby points. In order to determine if our embeddings capture these properties of related Pfam domains, we plot every Pfam domain that falls under the ten most common Pfam superfamilies (clans) in our self-supervised dataset: NADP Rossman (CL0063), P-loop NTPase (CL0023), Zn Beta Ribbon (CL0167), E-set (CL0159), HTH (CL0123), TPR (CL0020), PDDEXK (CL0236), MBB (CL0193), Beta propeller (CL0186), and OB (CL0021) [33].

**Fig 3.**
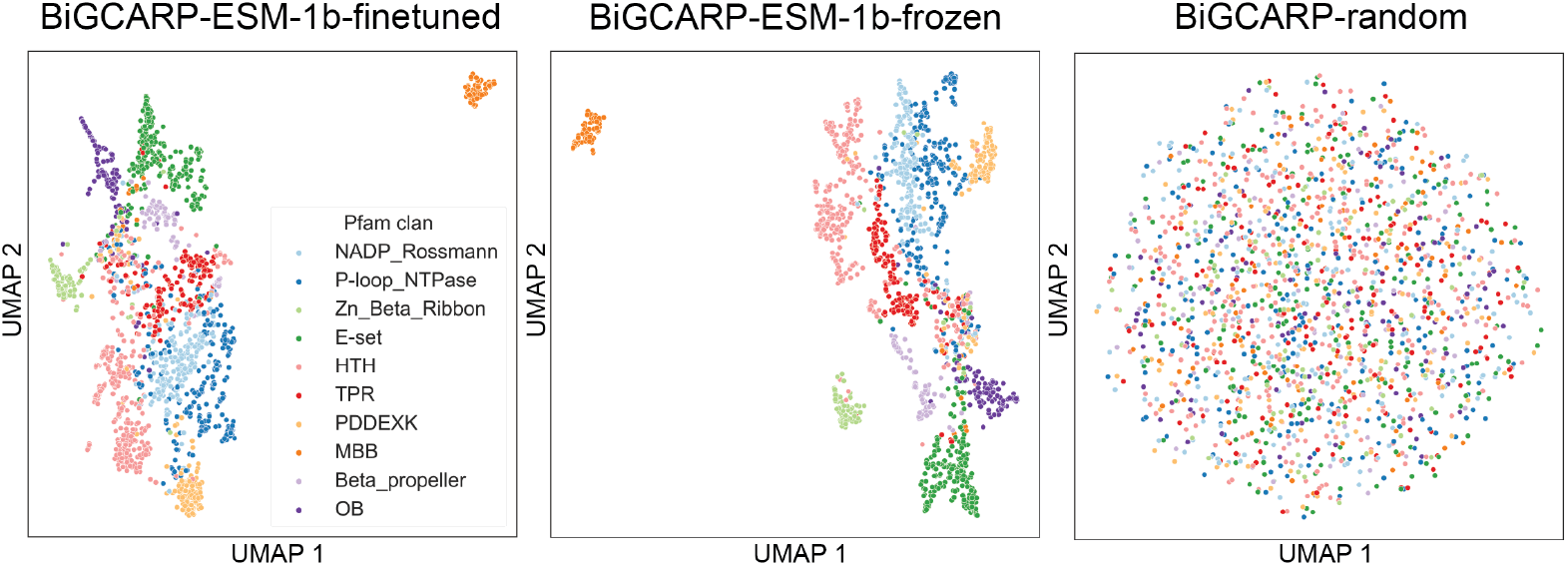
Relevant representations of Pfam domains are encoded in learned ESM-1b embeddings. Uniform manifold approximation and projection (UMAP) visualization of learned representations of Pfam domains from BiGCARP with different initial Pfam embeddings.

We find that initializing Pfam domain embeddings using ESM-1b improves the quality of the learned representations, as these embeddings take into account protein family amino-acid sequence. Fig. 3 indicates BiGCARP-ESM-1b and BiGCARP-ESM-1b frozen embeddings form clear clusters of structurally related Pfam domains, whereas using randomly initialized Pfam embeddings shows minimal interpretable information after self-supervised BGC training.

### BiGCARP captures meaningful patterns in BGCs

We next evaluated BiGCARP’s pretraining performance after self-supervised training (Table 1). We use the exponentiated cross entropy (ECE) metric for evaluating BiGCARP. This metric provides a measure for a model’s ability to narrow its prediction of a token from the set of options. An ideal model would have an ECE of 1, whereas a model choosing at random would have an ECE of the vocabulary size, which in our case is 19,550 for Pfam domains and 55 for BGC product classes. On our antiSMASH dataset test set, BiGCARP-ESM-1b-finetuned achieves the lowest ECE on the Pfam domains, while BiGCARP-ESM-1b-frozen achieves the lowest ECE on the product classes despite performing worse on domain ECE.

**Table 1.** Pretraining results, including the exponentiated cross entropy (ECE) metric on the pretraining test set and area under the receiver operating characteristic curve (AUROC) for BGC start locations and domains on the 9-genomes validation set.

|  |  | BiGCARP |  |  |
| --- | --- | --- | --- | --- |
|  |  | ESM-1b-finetuned | ESM-1b-frozen | random |
| pretrain test set (ECE) | Pfam domain product class | **4.64**<br>1.50 | 4.99<br>**1.46** | 4.67<br>1.50 |
| 9-genomes (AUROC) | start domain | 0.720<br>**0.876** | 0.701<br>0.611 | **0.723**<br>0.856 |

In addition, using the 9-genomes validation set from DeepBGC [10], we evaluate whether BiGCARP can identify the start locations of BGCs and whether each domain is in a BGC without further supervised training. We append a mask token to the beginning of every window of 64 domains in the dataset and pass them through BiGCARP. Intuitively, if the window is the start of a BGC, the model’s BGC class prediction should have low entropy, and its reconstructions of the domains should be both low-entropy and have low cross-entropy with the original input domain. This scheme is shown in Fig. 1. We refer the reader to Materials and Methods for details about scoring start positions and BGC Pfam domains. As shown in Table 1, all three versions of BiGCARP can detect BGC start locations and whether domains are part of a BGC, with BiGCARP-ESM-1b-frozen performing worse on both tasks than the other two versions.

We then finetuned BiGCARP on the training dataset reported in DeepBGC v0.1.0 [10], which uses all BGC domain sequences from MiBIG (version 1.4) as positive BGC samples and 10128 negative examples for 100 epochs and choose the epoch with the lowest validation loss on the 9-genomes validation set for further testing (Methods). Table 2 shows domain-level classification performance using area under the receiver operating characteristic curve (AUROC) on the 9-genomes validation set and the 6-genomes test set from DeepBGC. Note that the DeepBGC results on 9-genomes are for cross-validation directly on 9-genomes. All three versions of BiGCARP outperform DeepBGC on the 6-genomes test set and 9-genomes validation set. However, self-supervised training did not improve performance on the 6-genomes test set for BiGCARP.

**Table 2.** Domain AUROC after supervised training on the DeepBGC training set.

| pretraining | 6-genomes | 9-genomes |
| --- | --- | --- |
| BiGCARP-ESM-1b-finetuned | **0.950** | **0.941** |
| BiGCARP-ESM-1b-frozen | 0.946 | 0.940 |
| BiGCARP-random | 0.943 | 0.936 |
| none | **0.950** | 0.937 |
| DeepBGC | 0.921 | 0.934 |

### BiGCARP predicts BGC product classes

In addition to detecting BGCs in microbial genomes, predicting their product classes would provide further aid in discovering new natural products. BiGCARP learns to predict a BGC’s product class from its Pfam sequence by reconstructing masked class tokens during self-supervised training (Fig. 1). During self-supervised training, we use the antiSMASH product classes. In order to compare BiGCARP’s performance to DeepBGC, we map antiSMASH product classes to those in the Minimum Information about a Biosynthetic Gene cluster (MIBiG) dataset used in DeepBGC [10, 34].

DeepBGC trains a random forest classifier on its embeddings to predict BGC product classes. In contrast, we simply append a mask token to the beginning of each BGC sequence and evaluate the model’s predictions for the identity of the mask, removing the need to train an additional model.

All three versions of BiGCARP out-perform DeepBGC on average across the product classes, and an ensemble of their predictions further improves accuracy, as shown in Table 3 and Table S1. BiGCARP-ensemble outperforms DeepBGC on four out of seven product classes. This is likely because the antiSMASH training set is approximately 100-times larger than MIBiG. Performance is generally similar for product classes that are well-represented in both datasets, with the largest gains coming in the “other” and alkaloid classes, which are under-represented in MIBiG. This underscores the importance and utility of training on a large and diverse BGC dataset. We note that DeepBGC is advantaged here by reporting 5-fold cross-validation results on MIBiG, while BiGCARP is not trained on any sequences from MIBiG.

**Table 3.** Product classification results (AUROC) on MIBiG.

|  | *n* MIBiG | *n* antiSMASH | BiGCARP-ensemble | DeepBGC |
| --- | --- | --- | --- | --- |
| polyketide | 644 | 21,679 | 0.898 | **0.903** |
| NRP | 433 | 14,655 | 0.898 | **0.907** |
| RiPP | 199 | 26,721 | **0.963** | 0.907 |
| saccharide | 179 | 144 | 0.773 | **0.811** |
| other | 154 | 26,909 | **0.763** | 0.583 |
| terpene | 120 | 16,049 | **0.869** | 0.824 |
| alkaloid | 39 | 331 | **0.820** | 0.607 |
| average |  |  | **0.855** | 0.792 |

## Discussion

Biosynthetic gene clusters (BGCs) are a promising source of natural products, but are difficult to discover, express, and characterize. Recent work in self-supervised deep learning has shown promise for modeling DNA, RNA, proteins, and glycans. We develop **Bi**osynthetic **G**ene **C**onvolutional Autoencoding **R**epresentations of **P**roteins (BiGCARP), a masked language model that learns representations of BGCs based on their Pfam domains, detects BGCs, and predicts their product classes. To our knowledge, this is the first work to use Pfam domains as tokens in a masked language model. We demonstrate that our model learns biologically-reasonable representations of Pfam domains. Representing BGCs as Pfam domains was a compromise between limiting the sequence length while having fine-grained sequence information. Models on the level of amino acid residues or the nucleotide sequence may be able to resolve more details at the cost of more computation. BiGCARP is a strong BGC detector even without seeing negative examples, and achieves state-of-the-art accuracy in product class prediction.

The BGC masked language model introduced here demonstrates promise for the expansion of BGC science and engineering. In natural language processing and protein engineering, masked language models are often fine-tuned on downstream tasks of interest. For BGCs, these downstream tasks could include predicting their expression conditions or the chemical structures of their products. Without fine-tuning, our models may be useful for detecting previously unknown BGCs in microbial genomes and predicting BGC product classes.

## Materials and methods

In this section, we elaborate on details of our self-supervised deep learning framework for detecting BGCs from bacterial genomes and classifying them into their natural product classes. The workflow is summarized in Figure 1, which consists of curating data, pretraining Pfam domain embeddings, training BiGCARP, and using BiGCARP to characterize BGCs.

### Data

#### Pretraining dataset curation

To curate our pretraining dataset, we ran antiSMASH (ANTIbiotics & Secondary Metabolite Analysis SHell) 2.0, a microbial genome mining tool for BGC identification and analysis [6], on a database of 6,200 full bacterial genomes and 18,576 bacterial draft genomes [7]. This led to 142,821 total BGCs spanning 55 classes identified for model development and evaluation. Our choice of representing BGCs as Pfam domains led to a vocabulary size of 19,500 unique Pfam domains collected from Pfam database versions 31 and 32 [33]. We also remove sequences from the self-supervised training and validation sets that contain substrings from or are substrings of sequences from the MIBiG, 9-genomes, and 6-genomes datasets from DeepBGC described below. This results in 127,294 BGCs in our pretraining dataset prior to data splitting. All datasets used can be found on Zenodo.

#### Pretraining data split for training and evaluation

Our training, validation, and test sets were produced from an 80/10/10 split of the total set. Note that random splitting of data is widely avoided in biological sequence modeling, since it leads to evaluation of overly simple generalization. For example, in protein modeling, one instead uses sequence-identity based splits as a proxy for evolutionary signal [35]. Proper splitting of BGCs is more complex, as evolution of BGCs is poorly understood. To reduce redundancy between the data splits, we ensured that no example in one set was a strict substring of an example in another set.

#### DeepBGC datasets for evaluating BGC detection and product classification

We evaluated the performance of our models and compared it to the DeepBGC model by testing its ability to detect BGCs within bacterial genomes and to predict their corresponding product classes. To do this, we utilized DeepBGC’s training set with 617 positive and 10128 negative BGC samples to finetune our models [10]. We also used their 6-genomes and 9-genomes datasets to perform supervised domain classification tasks. For BGC product classification, we used DeepBGC’s MIBiG dataset, which contains 1406 BGCs. Our mapping from antiSMASH product types to common MIBiG compound classes can be found on Zenodo.

### Embeddings of Pfam domains with ESM-1b

We represent each Pfam as a vector. To do this, we take the first sequence in the alignment for a Pfam, then use ESM-1b [12], a protein masked language model, to embed all amino acids of this sequence. We averaged the embeddings over the full sequence, yielding a representation vector of size 1280. By obtaining pretrained embeddings of Pfam domains with ESM-1b, our model takes into account sequence details. To explore whether pretrained Pfam domain embeddings show improvement on the quality of Pfam domain representations, we use three different initial Pfam embeddings for BiGCARP: ESM-1b embeddings finetuned, ESM-1b embeddings frozen, and randomly initialized embeddings updated throughout training. ESM-1b-finetuned and ESM-1b-frozen have the same initialization at the start of self-supervised training. All other model weights were randomly initialized.

### BiGCARP architecture and training

We train BiGCARP using the masked language model objective described in [31]. We prepend a token representing the antiSMASH BGC class to each BGC sequence. Each sequence is then corrupted by changing some tokens to a special mask token or another Pfam domain token, and the model is tasked with reconstructing the original sequence. Specifically, 15% of tokens from each sequence are randomly selected for supervision during each training step. For those 15% of tokens, 80% are replaced by the mask token, 10% are replaced by a randomly-chosen Pfam domain token, and 10% remain unchanged. The model is trained to minimize the batch average cross entropy loss between its predictions for the selected tokens and the true tokens at those locations.

BiGCARP is a dilated 1D-convolutional neural network masked language model based on ByteNet [32] and CARP [30]. The input is a sequence of Pfam domains represented by 1280-dimensional vectors. Model hyperparameters include the following: kernel width of 3, maximum dilation of 128, 32 layers, and a hidden dimension of 256 for a total of 34 million parameters. Training parameters include the following: batch size of 64, Adam optimizer with a learning rate of 10^−4^, and mixed precision training using PyTorch [36] and NVIDIA Apex. Each version of BiGCARP was trained on one 32GB NVIDIA V100 GPU for 300 epochs. The epoch with the lowest validation loss was selected for downstream experiments. Model weights and datasets are available on Zenodo; training code and code to run pretrained BiGCARP models is available on Github. We do not report replicates for results as that would require training each model from scratch multiple times.

### Evaluation on 9-genomes and 6-genomes

We use the intuition that the model should make more confident predictions when given BGC sequences than non-BGC sequences to predict BGC start locations and whether each domain is part of a BGC. For each bacterial genome, we prepend a mask token to each possible subsequence of 64 domains and pass the resulting sequences to BiGCARP. With the exception of domains at the beginning and end of the genome, each domain is thus scored 64 times. For each window, we calculate the entropy of the predictions for the prepended mask token (start entropy), the entropy for each of the 64 domains in the window (domain entropy), and the negative log-likelihood of each domain in the window (negative log-likelihood). We predict whether a domain is the start of a BGC using the start entropy of the window for which it is the first domain; positions with a lower start entropy are more likely to be BGC start locations. We predict whether each domain is part of a BGC using the average of the start entropies for every window in which it appears and its domain entropy and negative log-likelihood within each window in which it appears (a total of 64 *×* 3 values). Domains with lower scores are more likely to be within a BGC.

### Supervised training on DeepBGC training set

We follow the supervised training procedure described in DeepBGC. Using the positive BGC domain sequences from MiBIG (version 1.4) and 10128 negative BGC domain sequences from DeepBGC, at each epoch, we shuffle the sequences into a “genome” and then predict whether each domain is part of a BGC. We fine-tune the self-supervised versions of BiGCARP as well as a randomly-initialized version using the Adam optimizer and a learning rate of 10^*−*4^ with early stopping using supervised results on 9-genomes.

## Supporting information

**Table S1.** Product classification results (AUROC) for individual BiGCARPs on MIBiG.

|  | ESM-1b-finetuned | ESM-1b-frozen | random |
| --- | --- | --- | --- |
| polyketide | 0.885 | 0.883 | 0.898 |
| NRP | 0.886 | 0.881 | 0.887 |
| RiPP | 0.960 | 0.950 | 0.950 |
| saccharide | 0.772 | 0.752 | 0.726 |
| other | 0.754 | 0.733 | 0.750 |
| terpene | 0.854 | 0.762 | 0.848 |
| alkaloid | 0.763 | 0.817 | 0.807 |
| average | 0.839 | 0.825 | 0.837 |

## Acknowledgments

This research was conducted using computational resources and services at Microsoft. We thank David Prihoda for assistance with the DeepBGC validation and test datasets and Jackson Cahn for inspiring discussions on BGCs.

## Author Contributions

**Conceptualization:** CRM, NB, KKY

**Data Curation:** CRM, NB, KKY

**Formal Analysis:** CRM, NB, KKY

**Methodology:** CRM, NB, APA, LC, KKY

**Project Administration:** APA, LC, KKY

**Software:** CRM, KKY

**Supervision:** APA, LC, KKY

**Visualization:** CRM, KKY

**Writing: Original Draft Preparation:** CRM, NB, KKY

**Writing: Review & Editing:** APA, LC

## Notes

https://doi.org/10.5281/zenodo.6857704

https://github.com/microsoft/protein-sequence-models

https://github.com/microsoft/bigcarp

### Competing Interest Statement

The authors have declared no competing interest.

